# TauG-Guidance Of Neonatal Vocalizing

**DOI:** 10.1101/143156

**Authors:** David N Lee, Mateo Obregon, Jonathan Delafield-Butt

## Abstract

A theory of action control (General Tau Theory) is applied to analyzing the vocalizations of human neonates. A central aspect of the theory, which is supported by experimental evidence across various actions and species, is that the trajectories of competent skilled actions follow a particular temporal pattern, which is described by the mathematical function, tauG. It was found that the acoustic waveform of vocalizations of healthy, full-term babies followed the tauG pattern with high precision. We conclude that healthy full-term neonates can be born with the ability to tauG-guide their vocalizations.

## Introduction

Humans and animals sculpt vocal sounds by moving vocal articulators – lungs, vocal folds (syrinx in birds), larynx, velum, jaw, tongue and lips. Therefore, understanding how animals vocalize requires understanding how the vocal articulators move. That is, a theory of vocalizing needs to be expressed within a general theory of movement - just as Kepler’s laws of planetary motion are expressed within Newton’s general laws of motion.

In contrast, the classical analysis and description of vocal sound (and instrumental sound) is based on Fourier analysis of the acoustic pressure wave, which makes no reference to the movements that produce the sound. Fourier analysis of vocal sound is inadequate for other reasons too. A major shortcoming is that vocal sounds frequently contain abrupt transients (corresponding, for example, to consonants in speech or singing, which are highly informative to the listener), whereas, in principle, Fourier analysis only applies to a repetitive wave of infinite extent. To describe the highly informative abrupt transients in speech and singing various fudge factors (e.g., involving wavelet analysis) have been introduced into the Fourier analysis of vocal sound. The end result is an ad hoc theory of vocal sound, still with no explicit recognition of the core role played by the movements of the articulators.

The idea of basing a theory of vocal sound on Fourier analysis would appear to stem from Helmholtz’s resonant harp model of hearing - which today is the basis of the place theory of hearing. In the resonant harp model, the first-order analysis of an acoustic wave consists in breaking it down into its component frequencies (corresponding to the strings of the ‘harp’). However, as Berstein (1967) pointed out, if the resonant harp model were correct then the simplest sounds for humans to identify should be pure frequencies, corresponding to resonance of a single ‘string’. Next, in ascending order of difficulty, should be chords, corresponding to resonance of a set of harmonically related strings. Finally should come speech with all its acoustic complexities, including rapid transients. The problem with the resonant harp theory of human hearing, or related theories, is that in natural hearing the order of difficulty in recognising sounds is completely the reverse of that predicted by the resonant harp model.

In short, the fundamental shortcoming of the classical Fourier analysis approach to analyzing speech and singing (and instrumental sound) is that it treats the sound as a disembodied phenomenon, instead of what it clearly is - the result of synergistic control of sets of muscles by the nervous system.

## Sculpting vocal sound

Sculpting vocal sound means sculpting the form of the acoustic pressure wave that emerges from the mouth. This pressure wave is the stimulus for hearing. The wave is sculpted both at the level of the elementary acoustic wavelets (from minimum to maximum to minimum pressure) and at the level of sequences of wavelets, which constitute phrases in speech and music. The elementary acoustic wavelets are sculpted by the respiratory muscles in the diaphragm and chest wall, acting in concert with the tension-regulating muscles of the vocal folds (syrinx in birds). This basic sculpting is then refined further up the vocal tract by movements of the larynx, pharynx, jaw, tongue, and lips. All these movements basically entail closing and opening *motion-gaps* (changing gaps between a current state and a goal state). The movement of the tongue up to and down from the palate when singing or talking is an example of a closing then opening motion-gap. The style in which the tongue-palate gap closes and opens affects the form of the vocal sound. General Tau Theory (Lee *et al*. 2009) predicts that in skilled speaking or singing (or indeed any skilled movement) the motion-gaps are closed and opened following the mathematical function *tauG*. The motion-gap is then said to be *tauG-guided*. The tauG-guidance equation governing a motion-gap, *X*, is *tauX* = *k tauG*, where *tauX* is the tau function of *X*, which equals, at any time, the current value of *X* divided by the current rate of change of *X* (in symbols: *τ*_*X*_ = *kτ_G_*). *TauG* (*τ_G_*) is the *tau* of the motion-gap between an object (e.g., Newton’s apple or an animal’s body) falling with constant gravitational acceleration from rest to the ground. 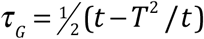, where time t runs from 0, at the start of the drop, to *T* at the end. Appropriately, the formula is derived from Newton’s laws of motion. There are three parameters, *k, A, T*, that determine how a *tauG-guided* motion-gap, *X*, opens or closes: *k* determines the *form* of the velocity profile of the motion-gap; *A* equals the *amplitude* of the motion-gap; *T* equals the *duration* of the motion-gap. If *k* is less than a half, the *X* motion-gap closes gently (with zero velocity at closure), the more so the lower the value of *k*. If *k* is greater than a half, the motion-gap closes forcefully (with positive velocity at closure), the more so the higher the value of *k*.

## Hypothesis

Based on General Tau Theory, our hypothesis is that, when vocalizing, the motion-gaps in the acoustic pressure wave, at both the wavelet and phrase levels, will be *tauG-guided*, to a degree depending on the level of vocalizing skill.

## Method

The study was carried out in the Neonatal Intensive Care Unit, Simpson Centre for Reproductive Health, the Royal Infirmary of Edinburgh, with the parents’ consent and before the babies were discharged from the hospital. The study was approved by the local Medical Ethics Committee.

### Babies

We recorded the vocalizations of 6 healthy term-birth babies when they were 24-72 hours old.

### Data recording

The baby lay supine on a bed with a parent at the foot of the bed, facing the baby. The parent was asked to converse with their baby. The baby’s vocalizations were recorded during three 5 minute periods, by holding a microphone (recording at 44,100 Hz) near the baby’s mouth. The three audio recordings were later divided into sections where the baby was vocalizing, and all these sections were then tauG-analyzed.

### Data analysis

Fig.1 and its caption explain the method of tauG-analyzing a baby’s vocalizations. The figure shows a representative sample from the present study. It is of a healthy full-term baby crying.

**Fig. 1.**
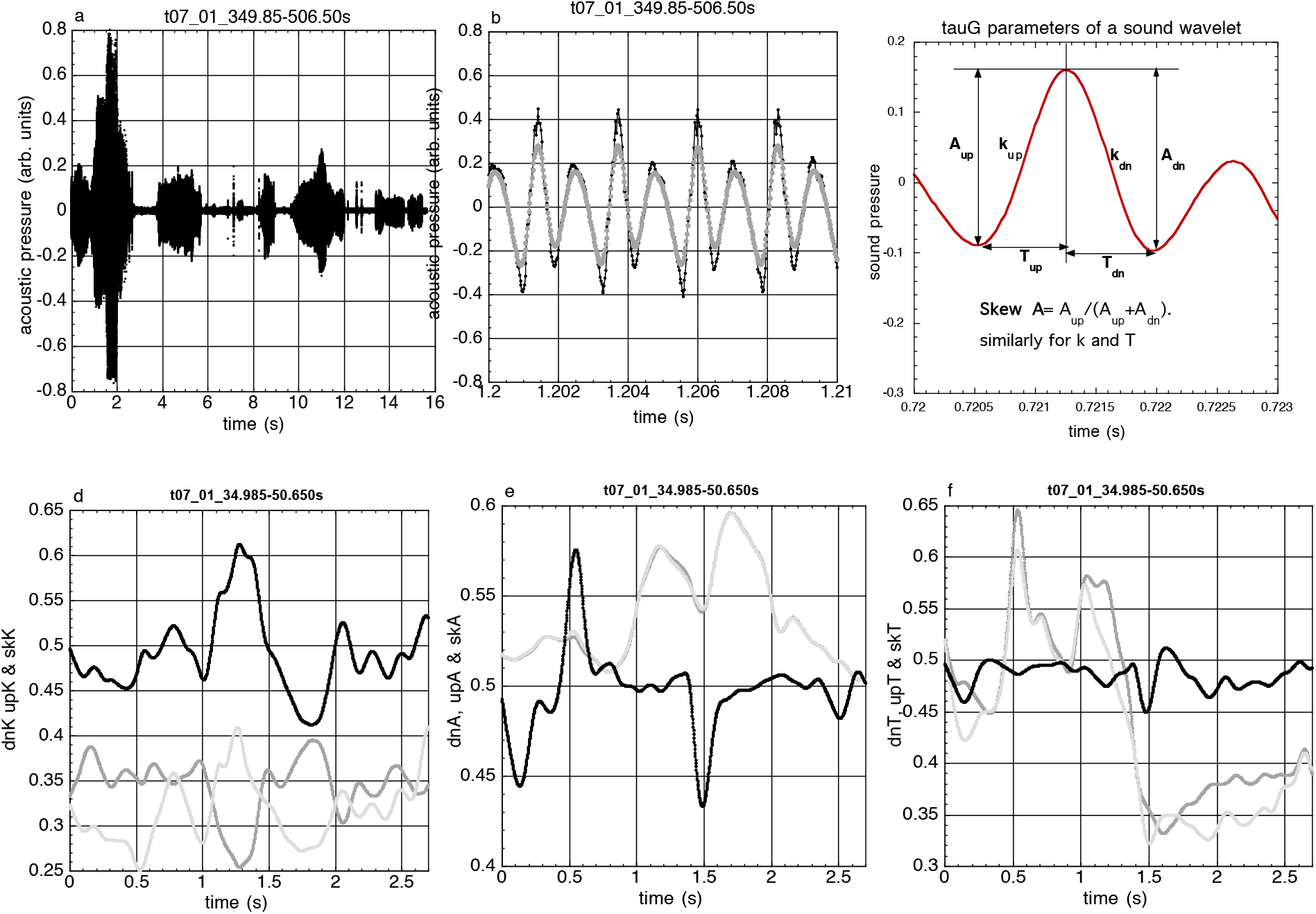
Illustrating tauG-analysis of a representative sample of a neonate’s cry (full-term baby t07). (a) The sound wave, recorded at 44,100 Hz. (b) a section of the sound wave (black) smoothed with a Gauss *σ* 9 filter (grey). (c) Schematic of a smoothed wavelet, showing the parameters computed by the tauG analysis program: *k_up_* and *k_down_* are the ‘*shapes*’ of the half-wavelets. These parameters are the coupling constants, *k*, in the *tauG-guidance* equation *τ_p_* = *kτ_G_*, where *P* is the pressure motion-gap to the peak or trough, respectively, of a wavelet. *A_up_, A_down_* are the *amplitudes* of the half-wavelets; (*A_up_*+*A_down_*)/2 approximates loudness. *T_up_* and *T_down_* are the *durations* of the half wavelets; 1/(*T_up_*+*T_down_*) approximates pitch. *k_skew_, A_skew_, T_skew_* measure the up/down asymmetry of the *k, A, T* parameters. *k_skew_ k_up_*/(*k_up_*+*k_down_*). Similarly for *A_skew_, T_skew_*. If symmetric, *skew* = 0.5. (d) The *k_skew_* (black), *k_up_* (light grey) and *k_down_* (middle grey) profiles during the initial section of the cry, 0 < t < 2.7 s. (e) and (f). Similarly for the parameters *A and T*.

## Results

Fig. 2 shows the mean results of a tauG-analysis of all five recorded cries of baby t07 (the same as illustrated in Fig. 1). The mean percentage of tauG-guidance (the measure of the degree of tauG-guidance) was high for both the up (black) and down (grey) sections all the profiles, particularly the phrase profiles. The other five of the six full-term babies studied showed similar results.

**Fig. 2.**
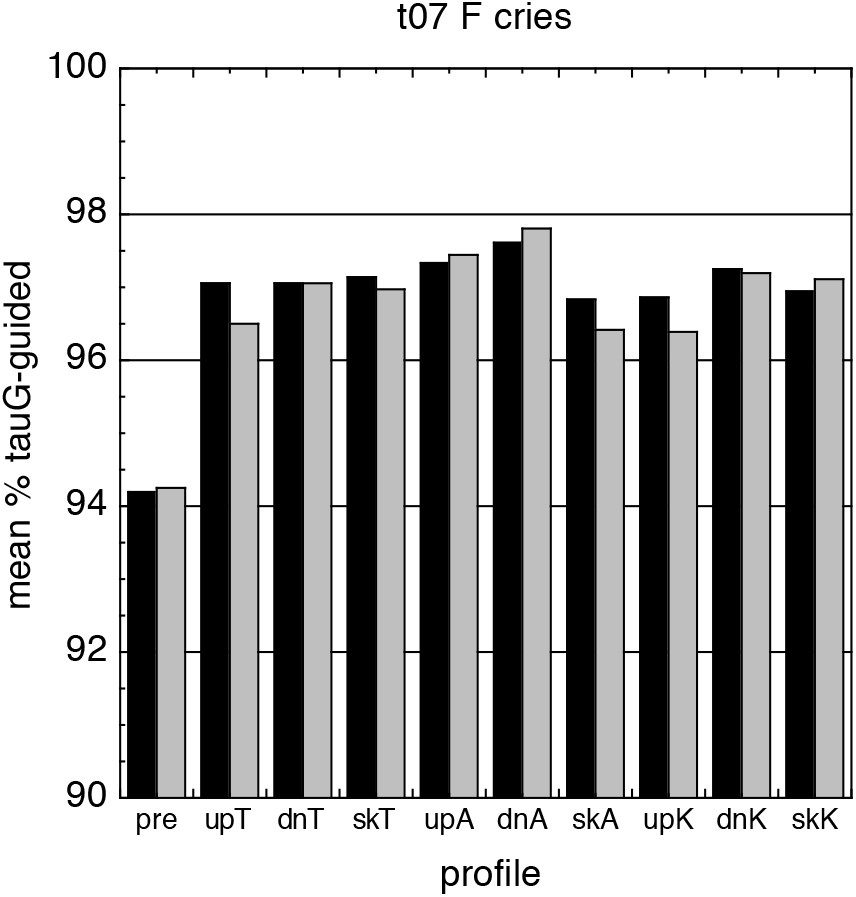
Mean percentage tauG-guidance of the cries of full-term neonate t07 (c.f. Fig. 1). Profiles are: pre: the sound pressure in the wavelets; upT and dnT: the durations of the up and down sections of the sound wavelets; skT: the up/down asymmetry in T (skT = 0.5 means symmetry; skT < 0.5 means upT < dnT). skA: the up/down asymmetry in A (skA = 0.5 means symmetry; skA < 0.5 means upA < dnA). skK: the up/down asymmetry in K (skK = 0.5 means symmetry; skK < 0.5 means upK < dnK). Black bars: the up sections of the profiles. Grey bars: the down sections of the profiles.

## Discussion

Our results indicate that healthy full-term neonates can be born with the ability to tauG-guide the sound wave of their vocalizations, both at the individual wavelet level and at the phrase level. This is perhaps not too surprising because, in general, young babies have to communicate effectively to have their needs satisfied. Therefore the basic elements of vocal communication in babies need to be the same as, or at least in tune with, the basic elements of vocal communication of adults, in singing and speech. Our other studies (e.g. Schogler, Pepping & Lee, 2008) indicate that basic elements of the sounds of adult singing, instrumental music and speech are also tauG-guided sound wavelets and tauG-guided phrase profiles.

This raises important medical possibilities. A tauG-analysis of neonates vocalizations could reveal a basic neuromotor dysfunction, which might then be treated through exposing the baby to music that is particularly replete with tauG-guided sound elements, to help the baby’s nervous system develop the capability of generating tauG patterns for vocalizing and many other activities too. This approach could be particularly valuable for young babies, because their nervous systems are more plastic.

